# Three heads are better than one: Cooperative learning brains wire together when a consensus is reached

**DOI:** 10.1101/2021.11.23.469804

**Authors:** Yafeng Pan, Xiaojun Cheng, Yi Hu

## Abstract

Theories of human learning converge on the view that individuals working together learn better than do those working alone. Little is known, however, about the neural mechanisms of learning through cooperation. We addressed this research gap by leveraging functional near-infrared spectroscopy (fNIRS) to record the brain activity of triad members in a group simultaneously. Triads were instructed to analyze an ancient Chinese poem either cooperatively or independently. Four main findings emerged. First, we observed significant within-group neural synchronization (GNS) in the left superior temporal cortex, supramarginal gyrus, and postcentral gyrus during cooperative learning compared to independent learning. Second, the enhancement of GNS in triads was amplified when a consensus was reached (vs. elaboration or argument) during cooperative learning. Third, GNS was predictive of learning outcome at an early stage (156-170 s after learning was initiated). Fourth, social factors such as social closeness (e.g., how much learners liked one other) were reflected in GNS and co-varied with learning engagement. These results provide neurophysiological support for Piaget’s theory of cognitive development and favor the notion that successful learning through cooperation involves dynamic consensus building, which is captured in neural patterns shared across learners in a group.

**Significance Statement:** Converging evidence has shown that cooperative learning is more effective than independent learning. An influential pedagogical theory postulates that learners benefit from cooperation through different forms of cognitive elaboration, such as providing elaborated clarifications to others. Alternatively, Piaget’s theory of cognitive development posits that cooperation encourages learners with diverse opinions to reach a consensus during the learning process. Here, we report that unlike individuals who worked alone, the brains of students who worked cooperatively with one another became synchronized. This within-group neural synchronization (GNS) was magnified when learners built mutual consensuses. These findings suggest that successful cooperative learning involves dynamic consensus building, which is reflected in the interpersonal coordination of cerebral activity.

## 1. Introduction

### Three heads are better than one

This well-known proverb, which appears in John Heywood’s 1546 collection (Heywood, 1972), implies that it is easier for multiple people to solve a problem or complete a task cooperatively than an individual working on their own. Similarly, education researchers have found that in peer-to-peer learning, students working in groups collaborate with and teach each other by addressing arguments (misunderstandings), providing elaborations (clarifications), and forming consensuses (Dillenbourg, 1999; van der Linden et al., 2000; Laal and Ghodsi, 2012). Interestingly, recent research that combines education and neuroscience have shown that social processes that take place during learning can be tracked through temporally synchronized neural processes among a group of students (Dikker et al., 2017; Davidesco et al., 2021). For example, within-group neural synchronization (GNS) in students predicted class engagement and how well they liked each other (Dikker et al., 2017). Little is known, however, about whether and how this type of GNS is involved in cooperative learning.

Despite the growing body of research about the cognitive and neural mechanisms of cooperative learning, there are still gaps in our understanding of why cooperative learning is better than independent learning. Educationists have interpreted this question in various ways. On the one hand, the Cognitive Elaboration perspective (Slavin, 1980; O’Donnell and Dansereau, 1992) postulates that by means of elaboration (e.g., understanding the learning content and explaining it to others), students working together can acquire knowledge far more effectively than students who work alone. Those who provided elaborated explanations to others also gained the most from cooperative learning (Webb, 1989; O’Donnell and Dansereau, 1992). Because of this, it was believed that the cognitive benefits of cooperative learning might arise from the learner’s (re)organization of knowledge and engagement with conspecifics. On the other hand, inspired by Piaget’s theory of Cognitive Development (Piaget, 1964; Huitt and Hummel, 2003), researchers have also hypothesized that cognition disequilibrium (e.g., disagreement on a view) emerges during cooperative learning because of prior information or educational diversity. For instance, members in a group may hold diverse opinions about certain learning content, leading to an uncomfortable state of cognitive imbalance (Ormrod, J. E. et al., 2016). When students reach a consensus with their peers, they benefit from that cooperation. This practice involves understanding others’ opinions and modifying one’s own views (schema) in order to return to a state of equilibrium (Kibler, 2011).

The present study sought to provide a neurophysiological testbed for the Cognitive Elaboration and Cognitive Development theories in cooperative learning. We aimed to explore learning interactions among a group of students from a neurophysiological perspective and investigate whether and how those learning behaviors (e.g., elaboration, arguments, and consensus) might bias this neurophysiological process. In recent decades, we have witnessed exciting developments in educational neuroscience approaches. For example, Nozawa et al. (2021) established that functional near-infrared spectroscopy (fNIRS) could be used to record brain activity from group members during cooperative learning simultaneously (the so-called “fNIRS hyperscanning”; Cui et al., 2012). This approach has been successfully applied in the field of learning sciences and consistently demonstrated that brain activity is synchronized between learners (and instructors) in naturalistic educational settings (Holper et al., 2013; Pan et al., 2018, 2020, 2021a; Zheng et al., 2018).

Thus, this study adopted fNIRS hyperscanning to investigate the mechanistic features of cooperative learning (vs. independent learning). We anticipated that cooperative learning would induce stronger within-group neural synchronization (GNS) compared to independent learning. Based on Piaget’s theory of Cognitive Development, we also believed that this type of GNS would be magnified when a consensus was built or, according to the Cognitive Elaboration theory, when elaborated explanations were delivered. Furthermore, inspired by a series of recent studies (Dikker et al., 2017; Bevilacqua et al., 2019), we also explored whether or not synchronized brain activity among learners during cooperative learning could predict learning outcomes and reflect group dynamics (e.g., social closeness and learning engagement).

## 2. Materials and Methods

### 2.1. Ethics statement

The ethics committee of East China Normal University approved the study. The study procedure was carried out following the Declaration of Helsinki. All participants were paid, and they gave their written informed consent before the experiment.

### 2.2. Participants

Sixty female, right-handed university students (all adults, mean age = 21.62, SD = 2.82) participated in the study, forming 20 triads (three-person groups). All participants were right-handed with no history of neurologic or psychiatric disorders. Same-gender volunteers were recruited to control for the potential effect of gender on social learning (Cheng et al., 2015; Baker et al., 2016; Pan et al., 2020). The participants in our sample had not tried to analyze or memorize a poem within the past two years.

### 2.3. Task, materials, and procedures

We used a poem learning task, which involved mastering two Chinese ancient poems (i.e., *Bu Suan Zi · Ode to the Plum Blossom*, 卜算子·咏梅, and *Bu Suan Zi · Huangzhou Dinghui Courtyard Residence*, 卜算子·黄州定慧院寓居作; see **Table 1**). The material was chosen because these ancient poems are typically taught in Chinese schools and are thus appealing to university students. Each poem only has eight lines and can be learned within 10 minutes. These two poems use similar types of rhetoric and flowery language (e.g., anthropomorphism) and convey the same emotions (e.g., lonely, unswervingly, and/or solitary). These pieces were selected from a classic national standard textbook (*The Collection of 300 Song Poems*). Using two poems allowed us to provide different learning contents for the two within-group conditions (i.e., cooperative learning vs. independent learning) without repetition.

**Table 1.** Learning materials in the poem learning task.

|  |
| --- |
| Poem #1 |
| Title: *Bu Suan Zi · Ode to the Plum Blossom* (卜算子·咏梅) |
| Author: *You Lu* (作者: 陆游) |
| *Outside the post-house, beside the broken bridge* (驿外断桥边) |
| *Alone, deserted, a flower blooms* (寂寞开无主) |
| *Saddened by her solitude in the falling dusk* (已是黄昏独自愁) |
| *She is now assailed by wind and rain* (更著风和雨) |
| *She craves not Spring for herself alone* (无意苦争春) |
| *Let other flowers be envious* (一任群芳妒) |
| *Her petals may be ground in the mud* (零落成泥碾作尘) |
| *But her fragrance will endure* (只有香如故) |
| Poem #2 |
| Title: *Bu Suan Zi · Huangzhou Dinghui Courtyard Residence* (卜算子·黄州定慧院寓居作) |
| Author: *Shi Su* (作者: 苏轼) |
| *Hanging sparse tung tree with missing moon* (缺月挂疏桐) |
| *Leaking the first quiet* (漏断人初静) |
| *See Youren coming and going alone* (谁见幽人独往来) |
| *Longly and lonely* (缥缈孤鸿影) |
| *Startled but turned around* (惊起却回头) |
| *Hate nobody* (有恨无人省) |
| *Picking up all the cold branches and refusing to perch* (拣尽寒枝不肯栖) |
| *Lonely sandbar is cold* (寂寞沙洲冷) |

During the experiment, two digital video cameras (Sony, HDR-XR100, Sony Corporation, Tokyo, Japan) placed in opposite directions were used to record learning behaviors among participant groups (**Fig. 1A**). Participants’ brain activity was recorded simultaneously through fNIRS over left fronto-temporo-parietal cortices (**Fig. 1B**). The process consisted of two task blocks, each of which were split into a resting-state phase (180 s), a learning phase (480 s), and a report phase (120 s). The two blocks differed with respect to the learning phase (cooperative learning vs. independent learning, **Fig. 1C**). The inter-block interval was approximately one minute. The order of the block was counterbalanced. Each block was paired with one poem, and the poem order was also counterbalanced. During the resting-state phase, three participants in a group (sitting face-to-face in a triangle, **Fig. 1A**) were asked to relax and remain still. During the learning phase, the triads analyzed the poem either cooperatively (condition #1) or independently (condition #2). In the cooperative learning condition, the participants were asked to work together to discuss and address the following three sets of short answer questions: (1) *What are the important images* (*Yi Xiang*, in Chinese) *in this poem, and what kind of atmosphere was rendered?* (2) *What is the most important verse, and what kind of subjective thoughts and feelings did the author express?* (3) *Which verses use rhetoric, and what kind of rhetoric was used?* These three sets of questions appear as representative tests in the classic national standard textbook. The participants in the group were told that after an 8-minute discussion, they had to select a representative to report their collective answers. In the independent learning condition, participants addressed the above questions by themselves (no communication was allowed). Immediately following the learning phase, there was a report phase, during which one representative was selected to (in the case of cooperative learning) or all members took turns to (in the case of independent learning) report their answers.

**Fig. 1.**
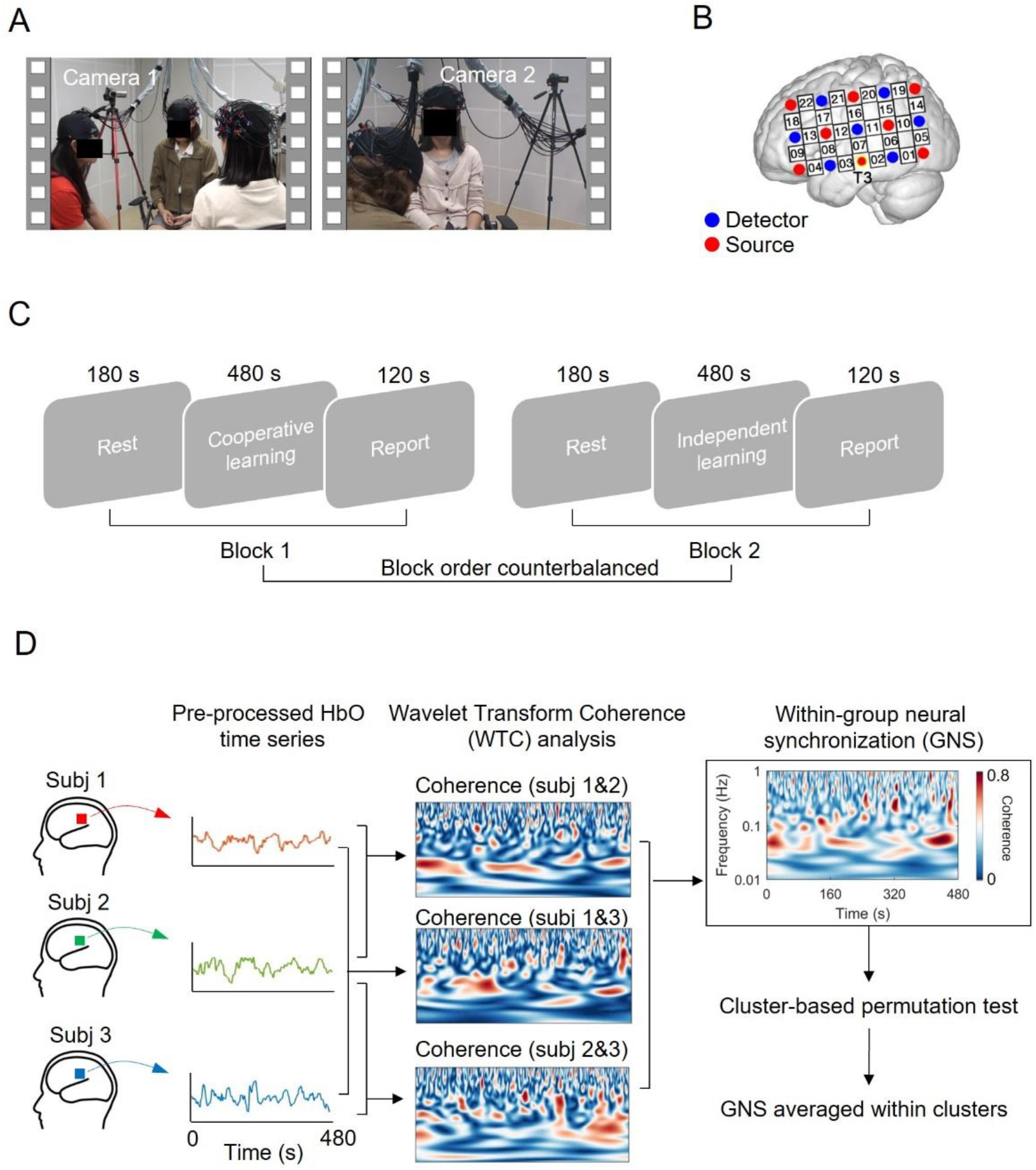
Experimental procedure, probe location, and within-group neural synchronization (GNS) computation. (A) During the experiment, three people within a group sat in a triangle. Two cameras were placed in opposite positions. Participants were asked to learn a poem in either a cooperative or an independent manner. (B) The fNIRS optode probe set was placed over the participant’s left fronto-temporo-parietal regions. (C) The participant group went through two task blocks (order counterbalanced), each of which included a 3-minute rest, an 8-minute learning period, and a 2-minute report. (D) Overview of the GNS analysis (based on a recently developed pipeline, Yang et al. 2020). Channel-wise raw oxy-hemoglobin (HbO) time series from each of the participants in a group were extracted and pre-processed. Interpersonal brain synchronization was estimated using Wavelet Transform Coherence (WTC) for two time series from each of the pairs (i.e., subject 1&2, 2&3, 1&3), resulting in three coherence matrices. GNS was computed based on the average of three coherence matrices. Data-driven cluster-based permutation tests were used to determine the frequencies and regions of interest. The data shown in the figure was based on a random triad.

### 2.4. Learning assessment

The answers that the learners provided for each question during the report phase constituted their learning performance. The performance of learners was scored by two separate raters who were blind to the experiment. For the cooperative learning condition, we took the score of the representative as the performance of the three-person group. For the independent learning condition, we averaged the score of the three members to calculate the overall performance of the group. Inter-rater reliability was calculated through the intra-class correlation (ICC) of scores (cooperative learning: ICC = 0.804, independent learning: ICC = 0.873). Rating scores were averaged across the two raters. The sum of the judgments made on all three questions for each triad served as an index of overall learning performance (maximum score = 15). A paired-sample *t*-test on learning performance was conducted to compare the difference between cooperative learning vs. independent learning.

### 2.5. fNIRS data acquisition

We used an ETG-7100 optical topography system (Hitachi Medical Corporation, Japan) to monitor and record brain data from triad members in a group simultaneously. The absorption of near-infrared light was measured with two wavelengths (695 and 830 nm). The sampling rate was 10 Hz. The modified Beer-Lambert law was used to estimate the oxy-hemoglobin (HbO) and deoxy-hemoglobin (HbR). One 3 × 5 optode probe set, which included eight sources and seven detectors forming 22 measurement channels, was used to cover each participant’s left fronto-temporo-parietal regions (**Fig. 1B**). These brain regions have been previously associated with social interactions in educational settings (Holper et al., 2013; Takeuchi et al., 2017; Pan et al., 2018, 2020, 2021a; Zheng et al., 2018). The distance between optodes was 3 cm. The middle optode for the lowest probe row of the patch was placed at T3, following the international 10-20 system. The middle probe set columns were placed along the sagittal reference curve. The Cz position in the 10-20 system, which lies midway in the two planes (i.e., nasion-inion and preauricular points), was used to examine and adjust the position of optodes within and across groups. A virtual registration approach was used to determine the correspondence between the NIRS channels and the measured points on the cortices (Singh et al., 2005; Tsuzuki et al., 2007).

### 2.6. fNIRS data analyses

#### 2.6.1. Pre-processing

Data collected during the learning phase (480 s) was extracted for subsequent analyses. A Correlation Based Signal Improvement (CBSI) approach was used to remove head motion artifacts (Cui et al., 2010). The CBSI approach reduced noise by maximizing the negative correlation (assumed to close to -1) between the HbO and HbR signals. It is defined by:

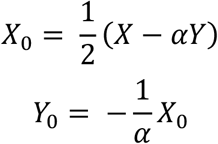

where *X, Y, X*_*0*_, *Y*_*0*_ denote raw HbO, raw HbR, corrected HbO, and corrected HbR signals, respectively, and α is the ratio of the standard deviation of raw HbO and raw HbR signals.

To remove global physiological noise (e.g., blood pressure, respiration) in fNIRS data, we used a wavelet-based denoising approach (Duan et al., 2018). Specifically, we automatically detected the time-frequency points that were contaminated by the global physiological noise using wavelet transform coherence. We then decomposed the fNIRS signal using the wavelet transform and suppressed the wavelet energy of the contaminated time-frequency points.

#### 2.6.2. Within-group neural synchronization (GNS)

This study only focused on the HbO concentration due to its higher signal-to-noise compared to HbR (Mahmoudzadeh et al., 2013), and because of its successful applications in fNIRS hyperscanning studies (Jiang et al., 2012, 2015; Dai et al., 2018; Hou et al., 2020). Also, note that our CBSI approach (see the *Pre-processing* section) maximally made corrected HbO and HbR signals negatively correlated; hence, parallel analyses on HbR returned analogous results. The pre-processed HbO time series was entered into the within-group neural synchronization (GNS) analysis (Yang et al. 2020, **Fig. 1D**). Following previous studies (Cui et al., 2012; Jiang et al., 2012), we used the wavelet transform coherence (WTC) method to estimate the cross-correlation between two HbO signals over homogeneous channels for each pair of the triad as a function of frequency and time (Grinsted et al., 2004). This analysis generated three WTC matrices for each triad (i.e., Coherence 1&2, Coherence 2&3, and Coherence 1&3). These three pairwise coherence matrices were then averaged as the group-level coherence (i.e., GNS) for each triad.

For futher analysis, GNS for the learning phase (480 s) was time-averaged and converted into Fisher-*z* statistics. To determine the frequencies and regions of interest, we employed a cluster-based permutation test (Maris and Oostenveld, 2007). The cluster-based permutation test is a nonparametric statistical test that allows for multiple comparisons for multi-channel and multi-frequency data (Maris and Oostenveld 2007, also see our recent application of this method in fNIRS hyperscanning, Zhu et al., 2021). The test consisted of five steps. First, we ran frequency-by-frequency (ranging from 0.01-1 Hz) and channel-by-channel (22 channels in total) paired-sample *t*-tests (cooperative learning vs. independent learning) on GNS. Note that we compared the two conditions directly (instead of relying on the task vs. rest comparison approach adopted by other studies) due to the fact that previous neuroimaging studies have shown that an active task condition could offer a better baseline than rest conditions (Stark and Squire, 2001; Reindl et al., 2018). Second, we identified channels (22 in total) and frequencies (80 in total, ranging from 0.01 to 1 Hz) that showed a significant *between-condition* effect (i.e., cooperative learning > independent learning, *P <* 0.05). Third, clusters with neighboring channels (N ≥ 2) and neighboring frequency bins (N ≥ 2) were formed. We computed the statistic for each cluster by summing all the *t* values. Fourth, we repeated steps one through three 1000 times using permutated data. The permutation was constructed by randomly paring one learner’s data in a group with the data of another learner from a different group (e.g., learner 1 in group 1 was re-paired with learner 2 in group 3 and learner 3 in group 7). GNS was estimated for the randomized groups in the same way that it was for the real groups. Finally, significant levels (*P <* 0.05) were calculated by comparing the obtained cluster statistic from the real group with 1000 renditions of randomized groups. This procedure identified two channel-frequency clusters that reached significance: Cluster 1 (frequencies 0.045-0.060 Hz, channels 11, 16, and 20) and Cluster 2 (frequencies 0.024-0.032 Hz, channels 9 and 13). These frequencies enabled removal of undesired signals in higher frequency bands, including those associated with Mayer waves (approx. 0.1 Hz), respiration (approx. 0.2–0.3 Hz), and cardiac pulsation (approx. 1 Hz). GNS results yield a *t* map for the left hemisphere. The *t* map was generated with a spatial interpolation method and rendered over a standard brain template (ICBM 152 nonlinear asymmetric template) using EasyTopo toolbox (Tian et al., 2013).

#### 2.6.3. Validation of GNS

GNS within each cluster was averaged. To validate that the above findings were not obtained by chance, we assessed the likelihood of finding significant GNS by using a *within-condition* permutation. Specifically, we re-analyzed the data after randomly pairing the participants for the cooperative learning condition (instead of cooperative learning vs. independent learning). This permutation was conducted 1000 times. Significant levels (*P <* 0.05) were obtained by comparing the cluster GNS from real groups with 1000 renditions of randomized groups. Only Cluster 1 remained after this validation. We thus constrained our subsequent analyses to Cluster 1.

#### 2.6.4. Linking GNS with learning behaviors

To better understand which behavior best contributed to GNS during cooperative learning, we sought to fuse the time course of GNS with video data. To do so, we down-sampled the GNS data to 1 Hz to obtain point-to-frame correspondence between the time series and video recordings. Two undergraduate students (S. Liu and Y. He) who were blind to the experiment were recruited to independently and manually code learning behaviors in the cooperative learning phase using the video-recording data. The two coders underwent training before the coding procedure. An educational expert (with thirty years of teaching experience) was invited to propose a coding schema and train the two coders to identify the cooperative learning behaviors correctly. Three types of learning behaviors were categorized (see **Fig. 3A** for example video frames): (1) *elaboration* (i.e., group members actively organized new knowledge and provided elaborated explanation to others); (2) *argument* (i.e., group members had disputes or disagreements over a certain concept or point of view); (3) *consensus*, (i.e., group members came to a general agreement over an opinion). Two coders coded every one second whether or not there were any of those learning behaviors. For all coding, inter-coder reliability was estimated according to the ICC (ICCs > 0.738). The two coders discussed the inconsistency of coding and came to an agreement. The coding procedure allowed us to compare GNS associated with different learning behaviors. A repeated-measure ANOVA on GNS was performed, with Type (elaboration vs. argument vs. consensus) serving as a within-subject factor. Post-hoc multiple comparisons were Bonferroni corrected.

To validate that GNS increased as a result of the occurrence of specific learning behaviors, we performed an event-related analysis. Specifically, we extracted averaged GNS in the time window of -6∼6 s relative to the onset of each learning behavior (elaboration vs. argument vs. consensus). The window size was set to 6 s because of the delay-to-peak effect in the fNIRS signal (Cui et al., 2009). To the extent that GNS was specific to a certain behavior, we would expect a rise of GNS associated with that behavior after (but not before) its onset. A repeated-measure ANOVA on GNS was performed with Type (elaboration vs. argument vs. consensus) and Time (before vs. after the onset of behavior) functioning as within-subject factors. Post-hoc multiple comparisons were Bonferroni corrected.

Moreover, inspired by a recent opinion paper (Cooney et al., 2020) that encouraged us to examine how a group conversation might contribute to GNS, we decided to conduct a second round of video coding by labelling the moment that each member in a triad was speaking. This practice allowed us to parse three additional types of conversation behaviors: (1) *airtime* (i.e., the total duration of speaking time); (2) *turn-taking* (i.e., the number of members taking turns to speak); and (3) *feedback* (i.e., a stream of “uh huh”s and “yeah”s to signal a member’s responsiveness to the speaker). The video coding was carried out by two graduate students (Y. Hua and X. Liu) who were not previously involved in the project and blinded to the experimental purposes. Inter-coder reliability was estimated for three types of communicative behaviors (ICC_airtime_ = 0.929, ICC_turn-taking_ = 0.842, ICC_feedback_ = 0.498), respectively. Given the low ICC for *feedback*, this type of behavior was excluded from the subsequent analysis. The two coders discussed the inconsistency in the coding and came to an agreement. Pearson correlational analyses were performed to test the potential contribution of group conversation (i.e., airtime and turn-taking) to GNS.

#### 2.6.5. The relationship between GNS and learning outcome

We first performed a Pearson correlation analysis on overall GNS (GNS averaged across the whole 480-s task duration) and learning outcome in the cooperative learning condition. We then repeated the above correlation analysis of GNS’s predictive power over learning outcome, but rather than averaging GNS across time, we performed more temporally precise computations. Specifically, we calculated the correlation between GNS and learning outcome for every second, resulting in 480 correlation coefficients. Pearson *r* values were fisher-*z* transformed to obtain a normal distribution. This analysis yielded a series of *p*-values that were corrected via the cluster-based permutation test (in the same way as previously mentioned, with exceptions that clusters were formed based on temporal adjacency, N ≥ 2 seconds, and cluster statistics were the sum of *r-* to*-z* values).

#### 2.6.6. The relationship between GNS and subjective measurements

After the experiment, we asked our participants to report their partner likeability and learning engagement based on a 9-point Likert. Ratings were averaged within each group. We then examined whether perceived closeness and engagement were reflected in GNS in the cooperative learning condition and whether they could predict learning outcome, using Pearson correlation analyses.

## 3. Results

### 3.1. GNS emerged as a result of cooperative learning

Overall, on the behavioral level, students learned the content better in the cooperative learning condition (mean ± standard error, 10.65 ± 0.34) than they did in the independent learning condition (9.24 ± 0.26, *t*_19_ = 3.89, *P <* 0.001).

On the brain level, the cluster-based permutation test revealed two significant frequency-channel clusters in which GNS during cooperative learning exceeded GNS during independent learning: Cluster 1 (frequencies 0.045-0.060 Hz, channels 11, 16, and 20; cooperative learning vs. independent learning, cluster statistic = 7.66, *P =* 0.01; **Fig. 2A**) and Cluster 2 (frequencies 0.024-0.032 Hz, channels 9 and 13; cluster statistic = 7.31, *P <* 0.001; **Fig. 2B**). Channels 11, 16, and 20 roughly cover the left superior temporal cortex, supramarginal gyrus, and postcentral gyrus, whereas channels 9 and 13 roughly cover the left inferior frontal cortex (Tzourio-Mazoyer et al. 2002).

**Fig. 2.**
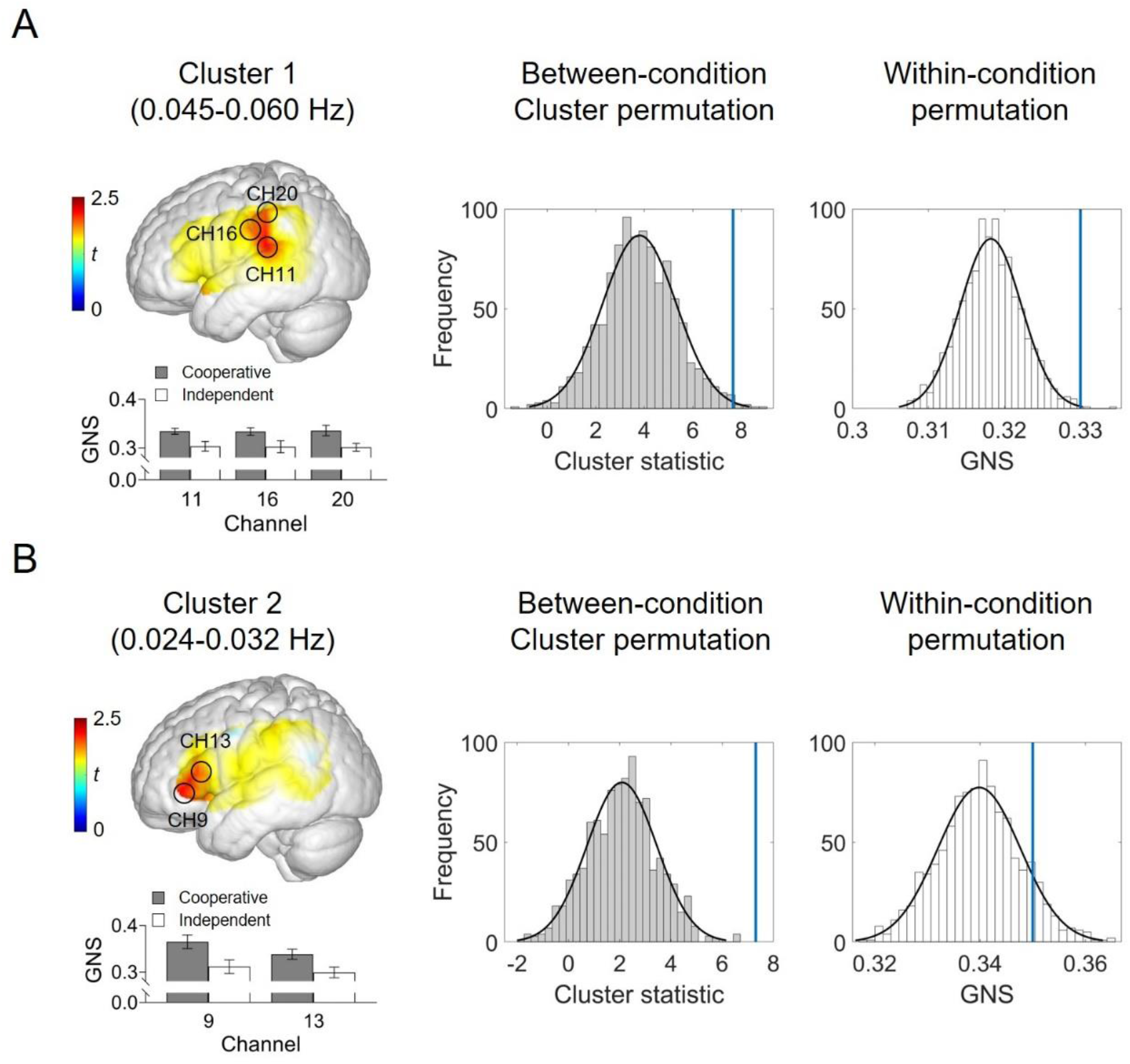
Within-group neural synchronization (GNS). Between-condition cluster-based permutation tests (cooperative learning vs. independent learning) revealed two significant clusters: (A) Cluster 1 (frequencies 0.045-0.060 Hz, channels 11, 16, and 20) and (B) Cluster 2 (frequencies 0.024-0.032 Hz, channels 9 and 13). Within-condition permutation tests (cooperative learning) verified that GNS over Cluster 1 (but not Cluster 2) did not emerge by chance. The blue lines indicate the positions of the cluster statistic or mean GNS for the original groups, relative to the distribution of the permutated data. In the bottom of brain models, the mean GNS for each condition and each channel within each cluster was displayed. The error bar indicates standard error.

A within-condition permutation analysis further confirmed that the patterns of GNS in the cooperative learning condition were specific to the interaction between real group members (pseudo members did not show higher GNS than real members), but only for GNS within Cluster 1 (*P <* 0.001, **Fig. 2A**, right panel). Cluster 2 showed similar levels of GNS between real and pseudo members (*P =* 0.13, **Fig. 2B**, right panel). Thus, in the remainder of the paper, we restrict our analyses to GNS within Cluster 1. GNS within Cluster 1 was averaged for subsequent analyses (cooperative learning vs. independent learning, 0.33 ± 0.03 vs. 0.30 ± 0.03).

### 3.2. GNS was more pronounced when a consensus was reached

To investigate which learning behaviors best contributed to GNS, we conducted a video coding analysis for each triad (**Fig. 3B**). The ANOVA revealed a main effect of Type, *F*_2, 38_ = 7.02, *P* = 0.003, *η*_partial_^2^ = 0.27, indicating that *consensus* elicited stronger GNS than *argument* (*t*19 = 3.72, *P*Bonf = 0.003) or *elaboration* (*t*19 = 3.76, *P*Bonf = 0.003). There was no significant difference between *argument*-GNS and *elaboration*-GNS (*t*_19_= 0.64, *P*_Bonf_ = 0.99, **Fig. 3C**).

**Fig. 3.**
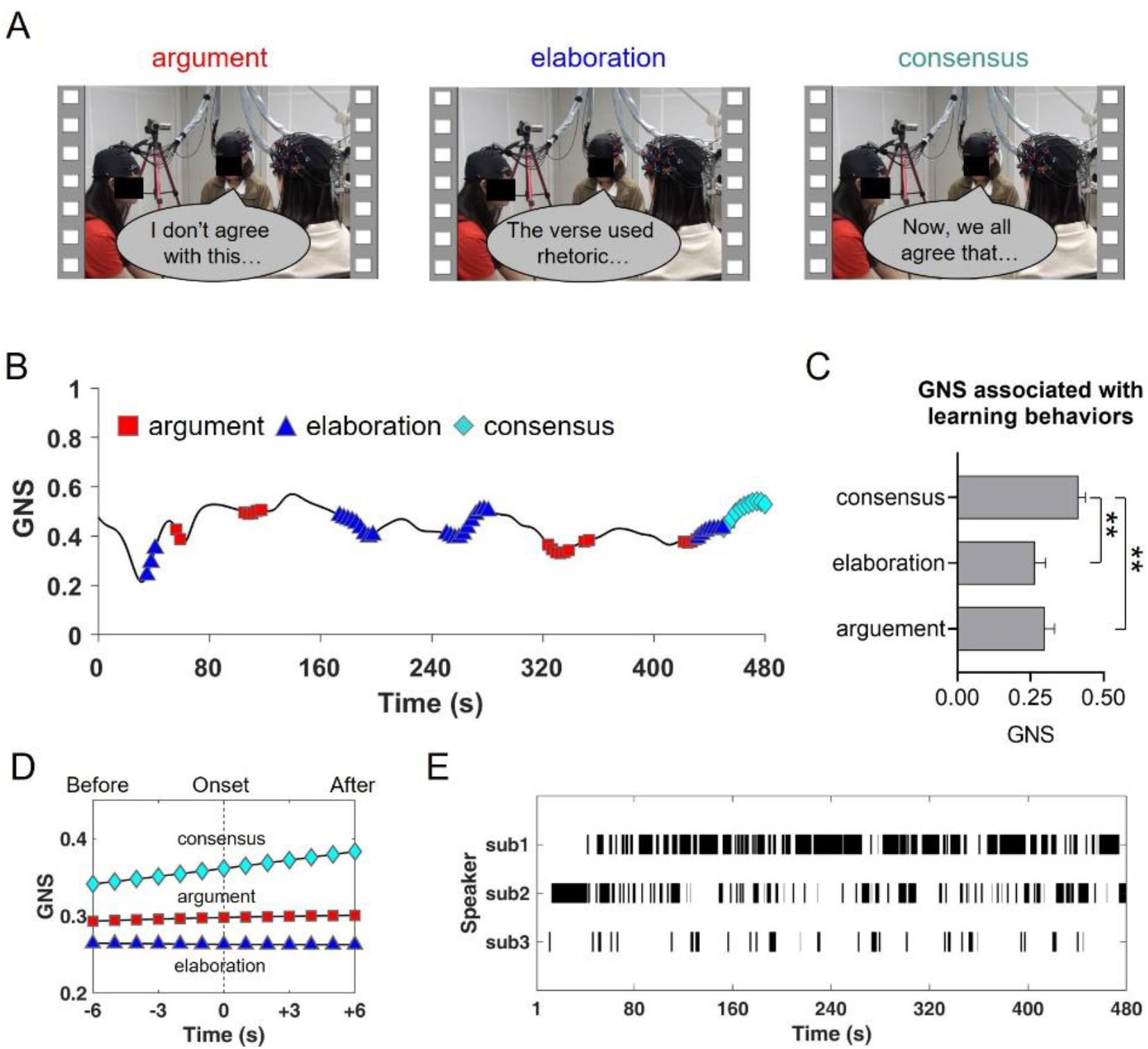
Within-group neural synchronization (GNS) as a function of time and learning behavior. (A) Example video frames coding learning behaviors. (B) Time course of GNS for a randomly chosen triad during cooperative learning. (C) GNS associated with consensus was stronger than that associated with argument or elaboration. The error bar indicates standard error. ** Bonferroni-corrected *P* < 0.01. (D) GNS associated with different learning behaviors in the time window of -6∼6 s relative to the onset of behaviors. (E) Moment-to-moment coding of group conversation for a randomly chosen triad during cooperative learning. The black vertical lines denote the moment when a member was speaking.

The event-related analysis on GNS in the time window of -6∼6 s relative to the onset of behavior revealed a main effect of Time (*F*_1, 38_ = 4.96, *P* = 0.04, *η*_partial_^2^ = 0.21) and significant interaction of Time × Type (*F*_2, 38_ = 8.89, *P* < 0.001, *η*_partial_ ^2^ = 0.32). Post-hoc analysis showed that *consensus* (but not other behaviors) elicited stronger GNS after rather than before (0.37 ± 0.09 vs. 0.35 ± 0.08, *P*_Bonf_ = 0.003) its occurrence (**Fig. 3D**). The main effect of Type was marginally significant (*F*_2, 38_ = 2.87, *P* = 0.07). Combined, these findings indicate that cooperative learning brains synchronized during the consensus moment.

Importantly, the effects reported above cannot be interpreted in terms of *airtime* (i.e., the duration of speaking) and *turn-taking* (i.e., the number of turn-takings) during group interaction, which were computed based on moment-to-moment coding of conversation (**Fig. 3E**), as we found no significant correlation between *consensus*-GNS and *airtim****e*** (*r* = -0.12, *P* = 0.61) or *turn-taking* (*r* = 0.15, *P* = 0.53).

### 3.3. GNS predicted learning outcome 156-170 seconds after learning onset

Contrary to our expectations, there was no significant relationship between overall GNS (GNS averaged across the whole time course) and learning outcome (*r* = 0.22, *P =* 0.36). Yet, when we ran the Pearson correlation analyses second-by-second, we found that GNS could significantly predict learning outcome 156-170 seconds after the onset of the learning phase (*r*s > 0.48, *P*s < 0.05; cluster statistic = 7.83, *P =* 0.03, **Fig. 4**).

**Fig. 4.**
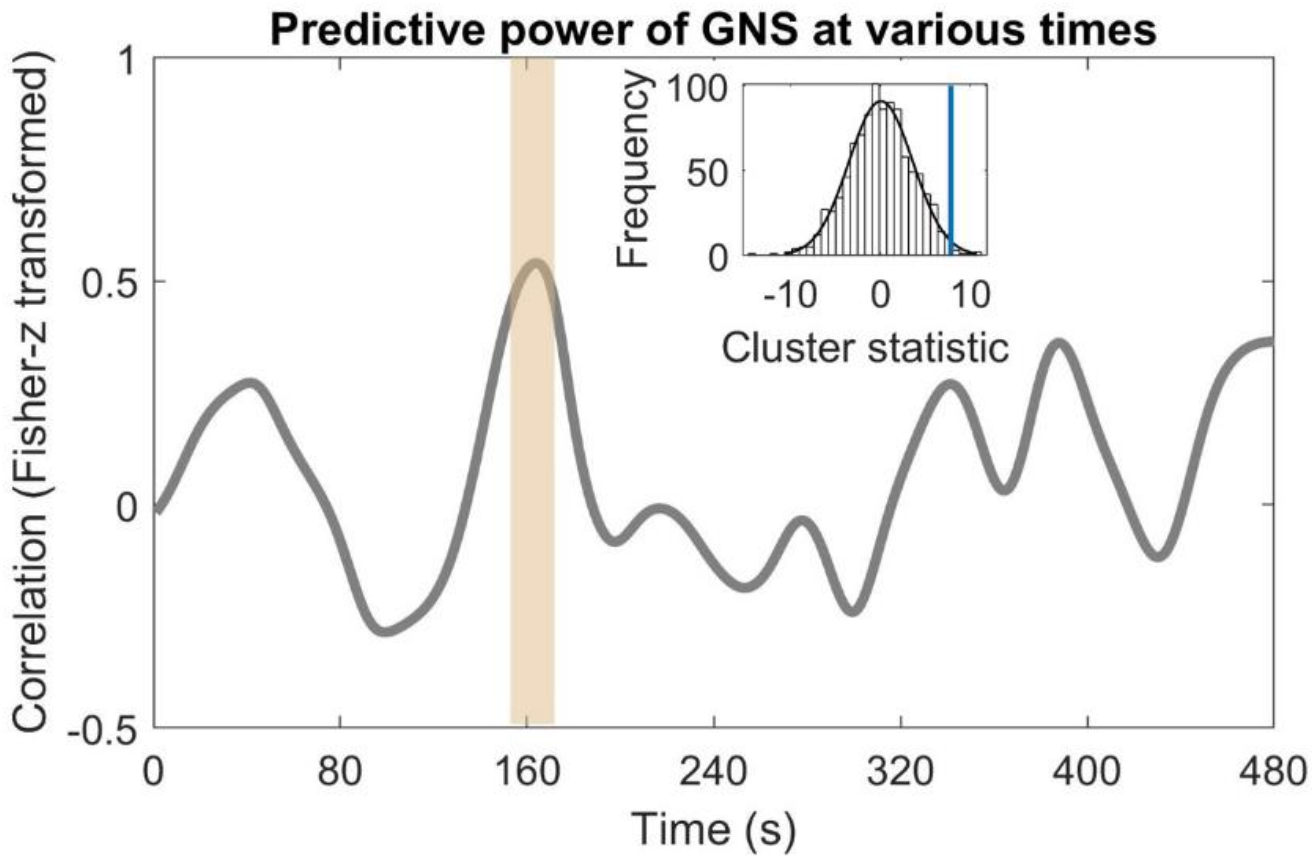
Predictive power of within-group neural synchronization (GNS) at various times. Time-varying Pearson correlation analysis revealed that GNS successfully predicted learning outcome at 156-170 s (indicated by a yellow vertical shadow) after the onset of the cooperative learning phase, confirmed by a cluster-based permutation test (the blue line indicates the position of cluster statistic for the original groups, relative to the distribution of the permutated data). Correlation coefficients were Fisher-*z* transformed.

### 3.4. Relationships between GNS, social closeness, and learning engagement

We observed a significant relationship between GNS and social closeness (i.e., how well learners like each other, *r* = 0.46, *P =* 0.04, **Fig. 5A**) post experiment. Social closeness also correlated with learners’ self-reported engagement (i.e., how learners engage during learning, *r* = 0.50, *P =* 0.02, **Fig. 5B**). Interestingly, engagement was associated with learning outcome (*r* = 0.45, *P =* 0.049, **Fig. 5C**).

**Fig. 5.**
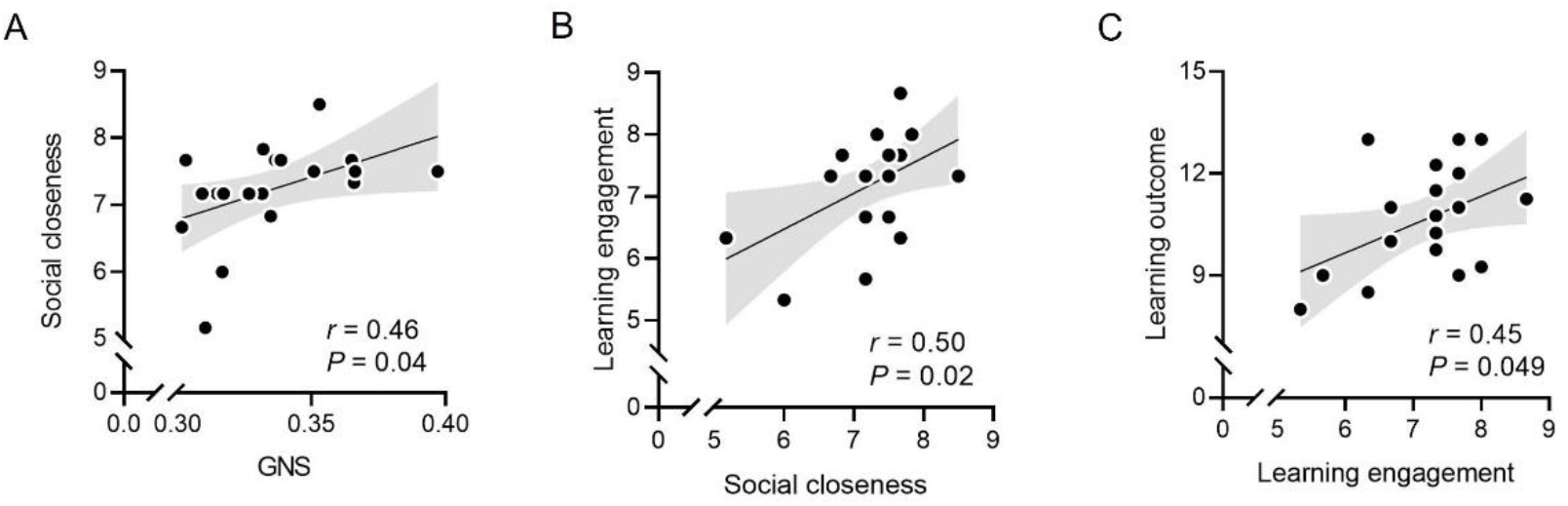
Relationships between within-group neural synchronization (GNS), social closeness, and learning engagement. (A) GNS correlated with social closeness, which was linked with learning engagement (B). (C) Learning engagement was associated with the learning outcome.

## 4.. Discussion

Cooperative group interaction can help improve the wisdom of crowds (Mercier and Claidière, 2021). In the present study, we used an fNIRS hyperscanning approach to investigate the mechanistic features of naturalistic cooperative learning. Both brain activity and behaviors were monitored and recorded simultaneously while triads were analyzing a poem in either a cooperative or independent manner. We coded learning behaviors (i.e., elaboration, argument, and consensus) based on video recordings, which provided a testbed for the Cognitive Elaboration and Cognitive Development theories. Our findings provided neurophysiological support for Piaget’s theory of Cognitive Development (Piaget, 1964; Huitt and Hummel, 2003), according to which cooperative learning facilitates cognitive developing by encouraging learners to reach a consensus with peers. We observed that cooperative learning elicited stronger within-group neural synchronization (GNS) in the left superior temporal cortex, supramarginal gyrus, and precentral gyrus relative to independent learning. Importantly, this type of GNS was pronounced when a triad reached a consensus during the learning process. Social factors such as closeness were reflected in GNS, which could predict learning outcome at an early stage (i.e., 156-170 s after the onset of learning).

Compared to independent learning, cooperative learning has the additional advantages of integrating verbal and nonverbal communication. This information integration triggered the alignment of neural processes among triads. The interpretation of GNS should be considered along with the function of coupled brain regions (i.e., superior temporal cortex, supramarginal gyrus, and precentral gyrus). The superior temporal cortex is an important hub for theory of mind (Baker et al., 2016) and has previously been associated with social perception and action observation (Thompson and Parasuraman, 2012). Previous studies have also shown that speaker-listener neural synchronization was stronger in the superior temporal cortex for more predictable speech (Dikker et al., 2014), suggesting that the superior temporal cortex plays a role in predictive coding in cooperative learning (Shamay-Tsoory et al., 2019). The supramarginal gyrus, like the superior temporal cortex, was recognized as being part of the temporo-parietal junction for theory of mind (Schurz et al., 2017). This region was widely observed in social animations. For example, movies with geometrical shapes that portrayed a social interaction could elicit activations within the supramarginal gyrus (Schurz et al., 2017). It is also involved in identifying nonverbal signals of other people during interaction (Reed and Caselli, 1994) and plays a vital role in empathy (albeit mostly in the right hemisphere, Silani et al. 2013). The postcentral gyrus, which is the location of the primary somatosensory cortex, is a critical region for the processing and integration of sensory and motor inputs. In this study, learners were not forbidden to use body language to facilitate cooperation. Taken together, GNS within these areas might imply that triads inferred one other’s understanding or mental states and integrated endogenous and exogenous information during cooperative learning. Certainly, developing a deeper understanding of the exact functional meanings of GNS within these regions could be the topic of future studies.

Notably, we also observed GNS in the inferior frontal cortex (i.e., Cluster 2). GNS over this area has been well established in face-to-face communication (Jiang et al., 2012). Our proposed nested permutation test (a cluster-based permutation between conditions, followed by a permutation test within the cooperative learning condition) permitted us to dissociate this component of verbal signal transmission from cooperative learning. Thus, it mitigated the concern that the GNS observed during cooperative learning (as opposed to independent learning) was simply due to verbal interactions in triads.

Based on our observation in the video coding analysis, the GNS effect was amplified when a consensus was reached. This finding echoes a recent study showing that group members’ neural activity became more aligned after they engaged in conversation with the aim of coming to a consensus about a movie clip’s narrative (Sievers et al., 2020). Our findings are better conceptualized when discussed alongside Piaget’s theory of Cognitive Development (Piaget, 1964; Huitt and Hummel, 2003). Based on this theory, learners benefit from forming agreements with peers during cooperative learning. Cognitive development is manifested through the process of transitioning from argument to consensus. In fact, we did observe that GNS associated with consensus exceeded GNS associated with argument. Consensus might require shared understanding among triads, which has been linked with neural synchronization in theory-of-mind-related regions between speakers and listeners (Stephens et al., 2010). Importantly, the effect cannot be simply attributed to the basic properties of face-to-face communication, such as airtime and turn-takings.

Like other studies (Dikker et al., 2017; Bevilacqua et al., 2019), we did not find a direct correlation between overall GNS (GNS averaged across the whole time course) and learning outcome. Time-varying correlation analysis, however, showed that GNS could positively predict learning performance at 156-170 s after the onset of the learning phase. While each group had its own discussion style and structure, preliminary consensus was reached around this time. The first argument tended to be resolved within the first three minutes after the learning process was initiated. Such time-varying correlation analysis provided more precise computations on a fine time scale compared to conventional correlation methods (Barnett and Cerf, 2017; Zhu et al., 2019).

Finally, GNS was found to correlate with social closeness (i.e., how much they liked each other). Learners who reported greater social closeness to conspecifics showed stronger GNS with conspecifics. This result replicated and extended previous evidence (Dikker et al., 2017; Bevilacqua et al., 2019) that social factors are important contributors for GNS during cooperative learning. Social closeness also correlated with learning engagement, which further predicted learning outcome. The relationships between these factors might suggest a “Brain (GNS) – Psychology (closeness) – Cognition (engagement) – Performance (learning outcome)” theoretical pathway implicated in the process of cooperative learning.

In summary, our study provided neurophysiological evidence for mechanistic features of cooperative learning. Students’ brains synchronized (wired together) during cooperative learning. Importantly, this effect magnified when triads reached a consensus, which is a finding that has potential benefits for cognifitive development. Future research projects could investigate whether exogenous manipulation of GNS through multi-brain stimulation (Novembre and Iannetti, 2021; Pan et al., 2021b) can causally enhance cooperative learning.

## Acknowledgment

We would like to thank Yingyao He, Shuo Liu, Yutong Hua, and Xiaotong Liu for their assistance in video coding. This work was supported by the National Natural Science Foundation of China (31872783 to Y.H. and 31800951 to X.C.) and the Shenzhen Basic Research Project (No. 20200810193259002) to X.C.

